# E2F deregulation compromises ER homeostasis by attenuating IRE1 activity

**DOI:** 10.1101/2025.06.02.657344

**Authors:** Arghya Das, Yining Li, Yiting Fan, Nam-Sung Moon

**Affiliations:** Department of Biology, McGill University, Montreal, Quebec, Canada

**Keywords:** Drosophila, Endocycle, Endoplasmic reticulum, E2F, Unfolded Protein Response, IRE1, Cytoplasmic DNA

## Abstract

The E2F family of transcription factors are key regulators of the cell cycle in all metazoans. While they are primarily known for their role in cell cycle progression, E2Fs also play broader roles in cellular physiology, including the maintenance of exocrine tissue homeostasis. However, the underlying mechanisms that render exocrine cells particularly sensitive to E2F deregulation remain poorly understood. The Drosophila larval salivary gland (SG), like its mammalian counterpart, is an exocrine tissue that produces large quantities of "glue proteins" in the endoplasmic reticulum (ER). Here, we show that E2F activity is important for the exocrine function of the Drosophila SG. The loss of *de2f1b*, an alternatively spliced isoform of Drosophila *E2F1*, leads to elevated DNA damage and accumulation of cytoplasmic DNA (cytoDNA) in the SGs. Surprisingly, we found that IRE1, a key sensor of the unfolded protein response, is required not only for ER homeostasis but also for preventing cytoDNA accumulation in the SG. Importantly, we found evidence demonstrating that IRE1 activity is attenuated in *de2f1b*-deficient SGs, contributing to both ER dysfunction and cytoDNA accumulation. Together, these findings reveal an unanticipated link between ER homeostasis and cytoDNA processing and offer mechanistic insight into why exocrine tissues are particularly vulnerable to E2F deregulation.

**Author Summary:** We discovered an unexpected consequence of E2F deregulation in ER homeostasis. The loss of *de2f1b* in the Drosophila salivary gland impedes physiological activation of an ER-resident protein, IRE1, a central component of the unfolded protein response. We show that IRE1 is required not only for ER homeostasis but also for preventing DNA accumulation in the cytoplasm, a phenotype observed in *de2f1b* deficient salivary glands. Our study reveals a previously uncharacterized role of IRE1-dependent ER function and offers new perspectives on why exocrine tissues are particularly vulnerable to the loss of E2Fs.

## Introduction

The E2F family of transcription factors are critical regulators of the cell cycle (1). Studies in metazoans have shown that E2Fs tightly control the expression of genes required for cell cycle progression. The essential role of E2Fs in the cell cycle is best illustrated by the fact that their activity is deregulated in nearly all cancer cells (2). Notably, while cell cycle regulation is the best-studied function of E2Fs, genetic studies across various model organisms have demonstrated that E2Fs affect a plethora of cellular physiology, such as apoptosis, metabolism, and cell-type specification (3–5). One of the understudied aspects of E2F biology is the vulnerability of exocrine cells to E2F deregulation. In mice, *E2f1* knockout leads to exocrine gland dysplasia in the salivary gland (SG) and pancreas, defects that become more pronounced with age (6). Moreover, *E2f1/E2f2* double-knockout mice and mice expressing hyperactive pRB, the primary inhibitor of E2Fs, develop diabetes due to defects in the pancreas (7–9). Gene expression profiling revealed that genes involved in exocrine and endocrine function are downregulated in *E2f1^-/-^/E2f2^-/-^*pancreases, suggesting that E2Fs promote and maintain terminal differentiation of pancreatic cells (8). Although the same study identified p53-dependent cell death as a contributing factor to pancreatic atrophy in *E2f1^-/-^/E2f2^-/-^* mice (10), the precise molecular mechanisms underlying the sensitivity of exocrine cells to E2F loss remain poorly understood. Intriguingly, acinar cells in the *E2f1^-/-^/E2f2^-/-^* pancreas become increasingly polyploid and accumulate DNA damage with age, correlating with the timing of pancreatic atrophy (7).

When the function of the endoplasmic reticulum (ER) is compromised due to the accumulation of misfolded or unfolded proteins, the unfolded protein response (UPR) is activated to restore and maintain ER homeostasis (11). The UPR consists of three branches: IRE1, PERK, and ATF6 signaling pathways. Among them, the IRE1 signaling pathway is conserved in all eukaryotes, ranging from yeast to humans (12). IRE1 is an ER transmembrane protein composed of the N-terminal ER luminal domain, a transmembrane domain, and the C-terminal cytoplasmic domain (13). Under normal unstressed conditions, its luminal domain interacts with an ER chaperone, Binding Immunoglobulin Protein (BiP), which prevents inappropriate activation of the IRE1 pathway. However, when unfolded proteins accumulate in the ER, BiP dissociates from IRE1, allowing IRE1 to oligomerize and activate its C-terminal domain that contains kinase and ribonuclease (RNase) activities. Of these, the RNase activity is critical for the IRE1-dependent UPR. The best-characterized substrate of IRE1 is X-box binding protein 1 (XBP1). The *xbp1* mRNA contains a short intronic sequence that disrupts the reading frame of its C-terminal domain. Upon ER stress, activated IRE1 excises this intervening sequence, leading to the translation of functional XBP1 proteins. In its active form, XBP1 promotes the expression of genes that enhance ER functions, such as ER chaperones and protein-folding enzymes (12). Notably, IRE1 is physiologically activated and essential for the development of exocrine tissues, which naturally have a high demand for ER-dependent protein synthesis (14). For instance, XBP1 deficiency in mice results in abnormalities specifically in secretory organs, such as the pancreas and the SG (15). The Drosophila E2F family is considered as a streamlined version of its mammalian counterpart. While mammals have eight E2F genes, Drosophila has only two: *de2f1* and *de2f2*, each representing distinct groups of mammalian E2Fs (1). Despite this simplicity, decades of research have demonstrated that E2F’s biological functions are well conserved between fruit flies and mice. Interestingly, *de2f1*, the only Drosophila E2F member capable of promoting cell cycle progression, undergoes alternative splicing (16). The difference between the canonical *de2f1* isoform, *de2f1a,* and the alternatively spliced form, *de2f1b,* is the inclusion of a microexon in *de2f1b*, coding 16 amino acids. Precise genomic deletion of the *de2f1b*-specific microexon demonstrated that *de2f1b* is specifically required in polyploid tissues such as the larval SG (16). Detailed analysis of the cell cycle revealed that a negative feedback loop that keeps Cyclin E/CDK2 activity in check is deregulated in *de2f1b*-deficient SGs (*de2f1b* SG), which leads to uncoordinated endoreplication (17).

Remarkably, the cells in the SG can have more than 1000 copies of their genome via endoreplication, an atypical cell cycle consisting of repeated G1 and S phases without intervening mitoses (18). This results in a Drosophila tissue with much higher DNA content per cell than a typical human diploid cell, making it an ideal system to study DNA biology. Another important aspect of the Drosophila SG is that, like its mammalian counterpart, it is an exocrine tissue. Starting at the mid-third instar larval stage, SG cells begin producing large quantities of “glue proteins” coded by salivary gland secretion (*Sgs*) genes (19). Indeed, the *Sgs* genes are associated with the polytene puffs, the chromosomal regions that are transcriptionally active in the polytene chromosomes (20). Throughout the late third instar larval stage, the glue proteins are synthesized in the ER and stored in secretory vesicles. Upon pupariation, stored glue proteins are secreted into the SG duct and, as the name suggests, they help to “glue” the puparium to solid surfaces during metamorphosis (21). Importantly, previous studies have shown that the IRE1 pathway is required in the larval SG, likely playing a crucial role in managing the high demand for glue protein synthesis (22).

In this study, we report that *de2f1b* SGs exhibit elevated levels of DNA damage and lead to the accumulation of cytoplasmic DNA (cytoDNA). Our investigation of the *de2f1b* SG phenotype demonstrates that physiological activation of the IRE1 pathway is attenuated in *de2f1b* SGs. Unexpectedly, IRE1-dependent ER function is required not only for expression of glue proteins but also for preventing cytoDNA accumulation, contributing, at least in part, to the defects observed in *de2f1b* SGs. Our findings explain why exocrine cells are particularly vulnerable to E2F deregulation and reveal a previously unappreciated link between ER homeostasis and cytoDNA processing.

## Results

Previous studies demonstrated that *de2f1b* mutant flies exhibit defects in endoreplicating tissues, including the Drosophila SG (16, 17). Given that E2F target genes are required for proper DNA synthesis, we asked if the polyploid genome of the *de2f1b* SG accumulates DNA damage and has an increased DNA repair activity. Notably, endoreplication in the SG naturally generates under-replicated regions, necessitating constant DNA repair to prevent the accumulation of free DNA ends (23, 24). Consistent with this idea, control SGs from mid third instar larvae (96-110 hrs. AEL, after egg laying) exhibited a basal level of 𝛾-H2Av foci, the Drosophila equivalent of 𝛾-H2AX foci (Figure 1A). Interestingly, we observed a higher number of 𝛾-H2Av foci in *de2f1b* SG nuclei, indicating an increased level of DNA damage and repair (Figure 1A). Since E2F targets also include genes required for DNA repair, we examined whether the genomic DNA of *de2f1b* SG is properly repaired via the terminal deoxynucleotidyl transferase dUTP nick-end (TUNEL) assay. While control SGs showed no detectable TUNEL signals, suggesting efficient DNA repair, numerous TUNEL-positive nuclei were observed in *de2f1b* SGs (Figure 1B). This result indicates that abnormal endoreplication in *de2f1b* SGs leads to heightened DNA damage that remains inadequately repaired during development.

**Figure 1.**
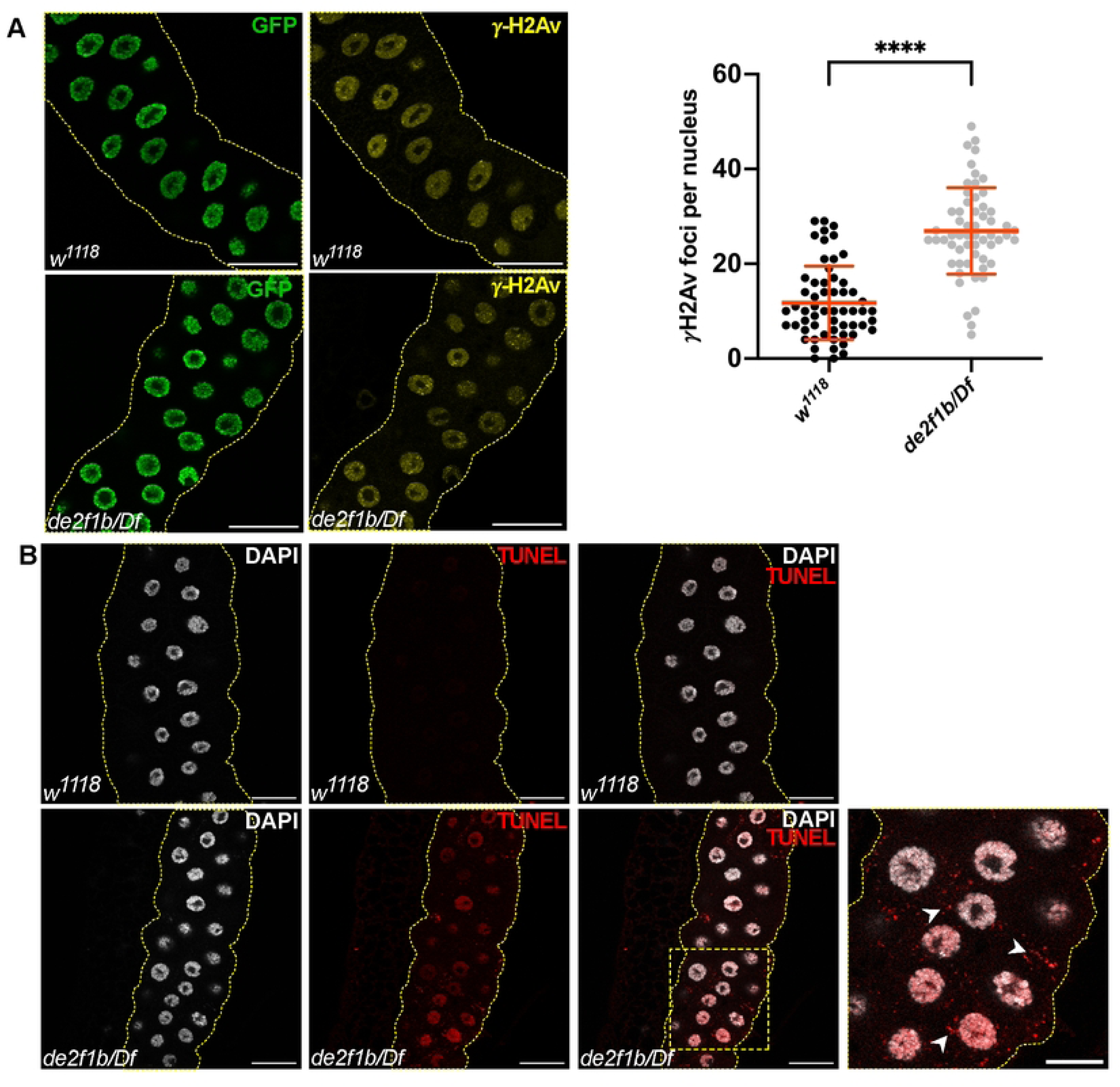
*de2f1b* salivary glands have an increased level of DNA repair and free DNA ends. (A) Control (*w^1118^*) and *de2f1b* (*de2f1b/Df*) salivary glands (SGs) expressing GFP-tagged Histone H2Av are stained with an antibody against γH2Av (yellow). Quantification of the number of the γH2Av foci normalized by the nucleus area is also shown. (****: *p* < 0.0001, scale bars: 50 μm). (B) The Terminal deoxynucleotidyl transferase dUTP nick end labeling (TUNEL) assays were performed with control and *de2f1b* SGs (red). Nuclei are also labeled with DAPI (white). A magnified view of the box area is also shown. White arrowheads mark cytoplasmic TUNEL signals (scale bars: 50 μm and 25 μm for the magnified images).

Strikingly, the TUNEL assay consistently produced cytoplasmic signals in *de2f1b* SGs, which were absent in controls (arrowheads in Figure 1B). To determine whether these cytoplasmic TUNEL signals represent the presence of cytoplasmic DNA (cytoDNA), we performed immunolabeling using an antibody that recognizes double-stranded DNA (anti-dsDNA). While strong cytoplasmic signals were absent in control, anti-dsDNA staining revealed robust signals in the cytoplasm of *de2f1b* SGs (Figure 2A). Notably, we frequently observed anti-dsDNA signals overlapping with weak yet discernible DAPI signals in the cytoplasm of *de2f1b* SGs (Figure 2A, lower panel). The anti-dsDNA was unable to produce a strong nuclear signal in the SG for unknown reasons. To rule out the possibility that the observed cytoDNA signals depict mitochondria, we examined the spatial correlation between anti-dsDNA signals and Mito-GFP, a mitochondrial marker comprising GFP fused with a mitochondrial localization signal (Figure 2B). While the anti-dsDNA was able to recognize mitochondria that were seen as cytoplasmic speckles (asterisks in Figure 2B), the prominent anti-dsDNA signals were clearly distinct from mitochondria (arrowheads in Figure 2B). Importantly, on no occasion did the cytoplasmic DAPI signals in *de2f1b* SGs overlap with Mito-GFP, indicating that the strong anti-dsDNA signals do not depict mitochondria.

**Figure 2.**
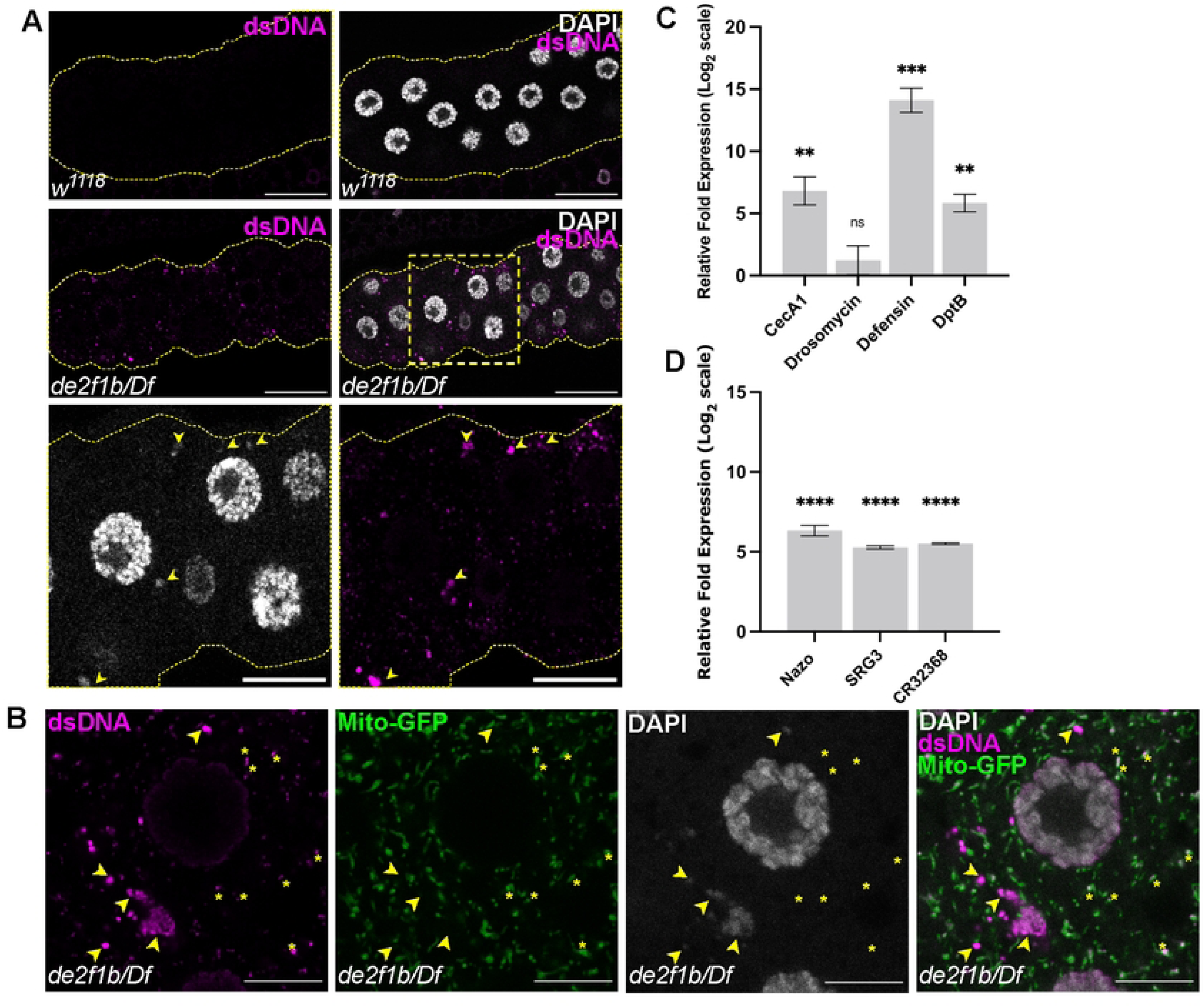
DNA accumulates in the cytoplasm of *de2f1b* SGs. (A) Control (*w^1118^*) and *de2f1b* SGs (*de2f1b/Df*) are stained with an antibody against double stranded DNA (anti-dsDNA, magenta) and DAPI (white). Lower panels show magnified views of the box area (scale bars: 50 μm and 20 μm for magnified images). Arrowheads mark anti-dsDNA signals that overlap with cytoplasmic DAPI. (B) *de2f1b* SGs expressing mitochondrially localized GFP (Mito-GFP, green) are stained with anti-dsDNA (magenta). Arrowheads mark anti-dsDNA signals that overlap with cytoplasmic DAPI and asterisks indicate anti-dsDNA signals that demark mitochondria (scale bars: 10 μm) (C) Antimicrobial peptide gene (AMP) expression levels were measured by RT-qPCR. Relative fold difference of indicated AMPs between control and *de2f1b* SGs are shown. (****: *p* < 0.0001, ***: *p* < 0.001, **: *p* < 0.01, *: *p* < 0.05). (D) The relative fold differences of the expression of previously identified STING-regulated genes between control and *de2f1b* SGs is determined by RT-qPCR. (****: *p* < 0.0001).

CytoDNA is a potent activator of the innate immune response (IIR) in mammals (25, 26). However, its role in the Drosophila IIR remains limited (27). For instance, while the mammalian cGAS (cyclic GMP-AMP synthase), which physically binds to cytoDNA and activates STING (Stimulator of Interferon Genes), has been well studied, the Drosophila cGAS ortholog that binds to cytoDNA has yet to be identified (28, 29). Regardless, we determined if the IIR is activated in *de2f1b* SGs by measuring the expression levels of several antimicrobial peptides (AMPs). In Drosophila, the IIR primarily engages the transcriptional activation of AMPs (30). As shown in Figure 2C, the expression levels of *Cecropin A1*, *Diptericin B*, and *Defensin* were greatly upregulated in *de2f1b* SGs. However, the expression level of *Drosomycin*, a widely studied Drosophila AMP, showed variable expression levels. Interestingly, RT-qPCR analysis also revealed that the expression of previously identified STING-regulated genes in Drosophila was highly induced in *de2f1b* SGs (31) (Figure 2D). Taken together, these data suggest that cytoDNA accumulates in *de2f1b* SGs and likely elicits the IIR.

To gain deeper molecular insights into the effects of the *de2f1b* mutation in the SG, we determined the gene expression profile by RNA sequencing (RNA-seq). While the best-studied function of dE2F1 is cell cycle regulation, ontology analysis unexpectedly revealed that genes involved in “protein processing in ER” were most significantly enriched among the downregulated genes in *de2f1b* SGs (Figure 3A and S1 Table). To validate the RNA-seq results, RT-qPCR on five genes within this category was performed. As shown in Figure 3B, all five genes were indeed downregulated in *de2f1b* SGs. This finding led us to further examine the ER morphology and its function. We used a GFP construct containing ER localization and retention signals, Bip-GFP- HDEL (ER:GFP), to visualize ER structures in the SG. In control SGs, a mesh-like ER network throughout the cytoplasm was observed (Figure 3C, upper panel). In contrast, *de2f1b* SGs displayed abnormal ER morphology, characterized by a less arborized and irregular ER network with varying ER:GFP intensities (Figure 3C lower panel). Notably, some cells were unusually small and appeared to completely lack the typical mesh-like ER network (arrowheads in Figure 3C). Since the glue proteins, the major proteins produced by the SG, are synthesized and folded in the ER, we investigated whether glue protein expression is affected in *de2f1b* SGs. To achieve this, we used a genomic construct where the carboxy-terminal region of a glue protein, SGS3, was replaced with the GFP coding sequence (SGS:GFP, Figure 3D). A previous study demonstrated that SGS:GFP expression follows a distinct spatiotemporal pattern, initially appearing at the distal tip of the SG and progressively expanding to the anterior cells (19). In control mid third instar larvae (96-110 hrs AEL), we observed a characteristic pattern where SGS:GFP is expressed in cells at the distal half of the SG while cells in the proximal half remain GFP-negative (Figure 3D upper panel). Notably, we noticed that the SGs expressing SGS:GFP have a discernible level of cytoDNA (see discussion). Nevertheless, this characteristic expression pattern of SGS:GFP is disrupted in *de2f1b* SGs (Figure 3D, upper panel). Cells lacking SGS:GFP expression appeared at random positions along the gland and these cells often contained a high level of cytoDNA (arrowheads in Figure 3D, lower panels). Together, our results indicate that the lack of dE2F1b causes ER abnormalities in the SG, causing failure in some cells to express glue proteins.

**Figure 3.**
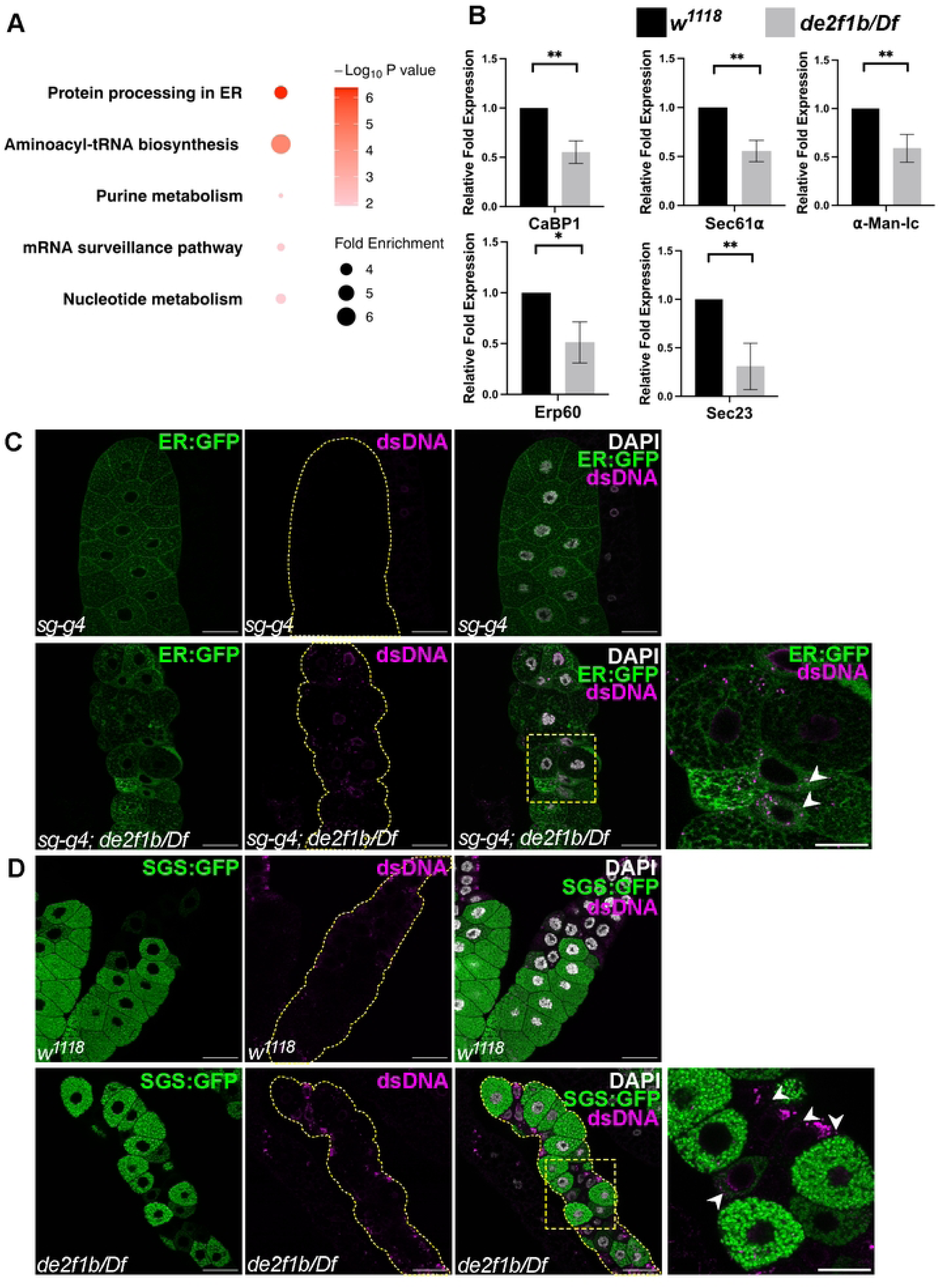
The endoplasmic reticulum (ER) homeostasis is deregulated in *de2f1b* SGs. (A) RNA-seq was performed to compare the gene expression profile between control and *de2f1b* SGs. Ontology analysis was performed with genes whose expressions are downregulated in *de2f1b* SGs. The top 5 biological processes identified from the ontology analysis are shown. The intensity of the red depicts the false discovery rate, and the size of the circles depicts the fold enrichment. (B) RT-qPCR was performed on five genes that are associated with “protein processing in ER” from ontology analysis. (***: *p* < 0.001, **: *p* < 0.01, *: *p* < 0.05). (C) A GFP construct tagged with ER localization and retention signals is used to visualize ER morphology (ER:GFP, green). Anti-dsDNA (magenta) was also used to determine the abundance of cytoDNA. Magnified view of the boxed area is also shown. (scale bars: 50 μm and 25 μm for magnified images) (D) A genomic construct, in which the coding region of the SGS3 gene is fused with the GFP sequence (SGS:GFP, green), is used to monitor the expression of “glue proteins” in control and *de2f1b* SGs. Anti-dsDNA (magenta) was also used to determine the abundance of cytoDNA. White arrowheads point to cells with a high level of cytoDNA. (scale bars: 50 μm and 25 μm for magnified images). FB: Fat body.

A closer examination of the genes associated with protein processing in ER (Figure 3A) revealed that a considerable number of their mammalian orthologs were previously identified as direct targets of XBP1, a transcription factor downstream of the IRE1 pathway (32). The XBP1- GFP construct is a commonly used sensor of the IRE1 activity where the GFP sequence is fused in frame with the C-terminal domain of XBP1 (33). Consequently, the GFP sequence of XBP1- GFP is translated only when IRE1 is active, and the intervening sequence preceding the C-terminal domain is removed (see introduction). As shown in Figure 4A, XBP1-GFP expression in control SGs resulted in clear GFP signals (Figure 4A, upper panel). This indicates that the Drosophila SG activates the IRE1 pathway during development, likely to cope with the high demand of ER-dependent glue protein synthesis (22). Interestingly, XBP1-GFP signals were overall weaker in *de2f1b* SGs than in the control and almost absent in some cells, indicating that the physiological activation of the IRE1 pathway is attenuated (Figure 4A). We next asked if the genes associated with protein expression in ER identified by RNA-seq (Figure 3B) are regulated by the IRE1 pathway. Indeed, the five genes downregulated in *de2f1b* SGs also showed reduced expression when either *ire1* or *xbp1* is depleted in the SG (Figure 4B). These results suggest that physiological activation of the IRE1 pathway is attenuated by *de2f1b* mutations, which likely contributes to the gene expression profile of the *de2f1b* SG.

**Figure 4.**
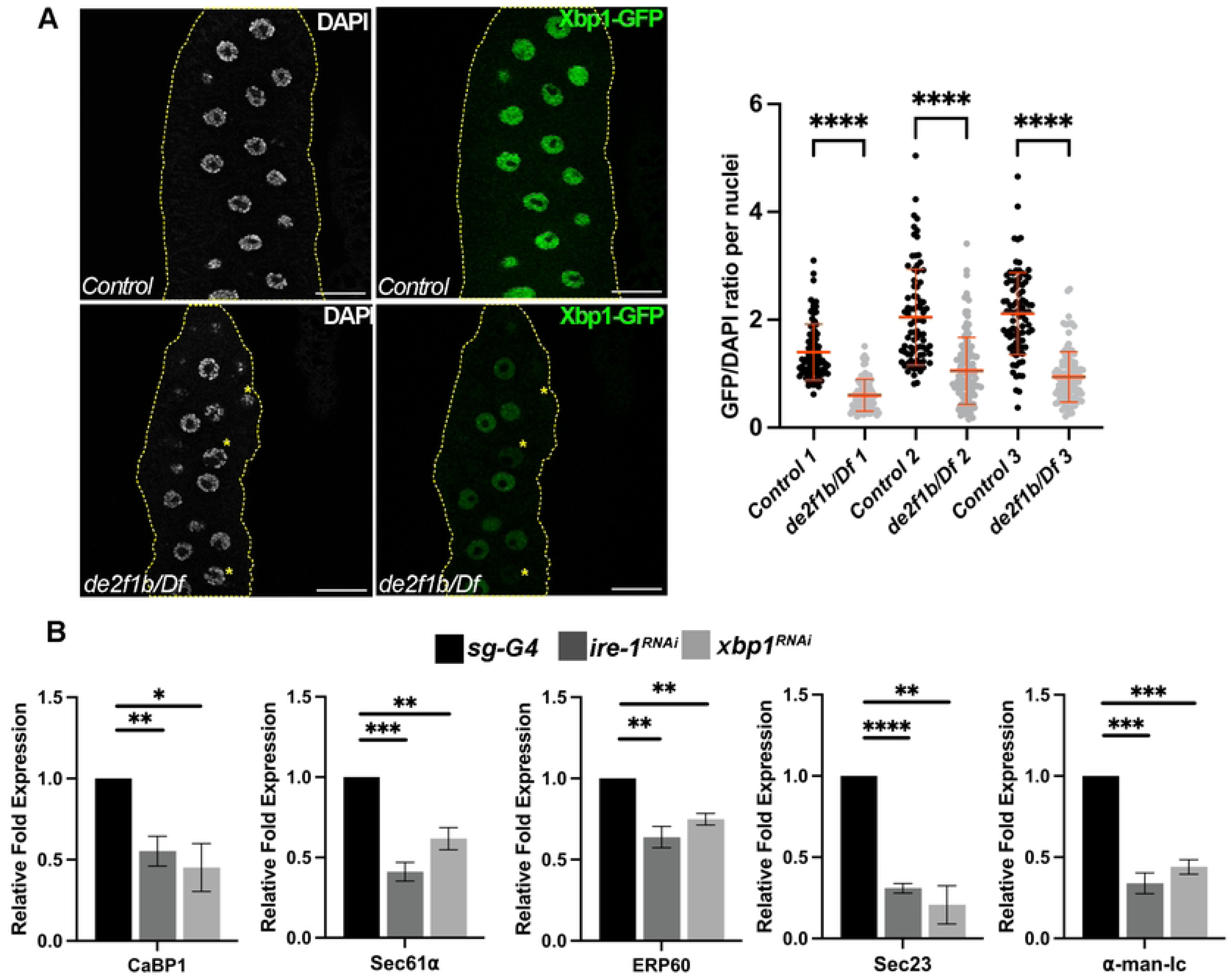
The IRE1 branch of the unfolded protein response (UPR) signaling pathway is deregulated in *de2f1b* SGs. (A) A construct, in which the GFP sequence is fused in frame with the C-terminal domain of XBP1 (XBP1-GFP), is used to monitor the IRE1-dependent XBP1 splicing (Scare bars: 50 μm). The dot plots show fluorescence intensity of XBP1-GFP per cell normalized by the DAPI signal in control and *de2f1b* SGs. Three independent experiments with matching controls are presented. (**** *p* < 0.0001) (B) Relative expression levels of the five genes whose expressions are downregulated in *de2f1b* SGs (Figure 3B) are determined in either *ire1* or *xbp1* depleted SGs. (****: *p* < 0.0001, ***: *p* < 0.001, **: *p* < 0.01, *: *p* < 0.05).

To investigate the role of the IRE1 pathway during SG development and to assess whether attenuated IRE1 activity contributes to the ER phenotype observed in *de2f1b* SGs, we depleted *ire1* or *xbp1* in wild-type SGs. ER:GFP revealed that IRE1 is required for proper ER development (Figure 5A). Depletion of *ire1* resulted in failure to form the mesh-like ER network observed in control, closely resembling the abnormal ER morphology observed in *de2f1b* SGs (Figure 3C). In contrast, *xbp1* depletion did not visibly affect ER morphology (Figure 5B). RT-qPCR confirmed the efficient depletion of *xbp1* by the RNAi construct (S1 Fig.) and the same *xbp1* RNAi construct was used to demonstrate XBP1-dependent expression of the five ER genes shown in Figure 4C. Therefore, this result suggests that an XBP1-independent function of IRE1, such as regulated IRE1-dependent decay (RIDD), may be critical for ER network formation in the SG (34, 35). Strikingly, *ire1* depletion in wild-type SGs was sufficient to produce cytoDNA, indicating that IRE1-dependent ER functions are likely required for preventing cytoDNA accumulation (Figure 5C). The functional importance of IRE1-dependent ER function during development is further demonstrated by the observation that *ire1*-depleted SGs failed to express high levels of SGS:GFP at the developmental stage when control SGs display a robust SGS:GFP expression (Figure D). Taken together, these results indicate that physiological activation of IRE1 is essential for proper ER development and that IRE1-dependent ER functions are necessary not only for glue protein expression but also for preventing cytoDNA accumulation. Furthermore, these results support the notion that attenuation of the IRE1 pathway likely contributes to the defects observed in *de2f1b* SGs.

**Figure 5.**
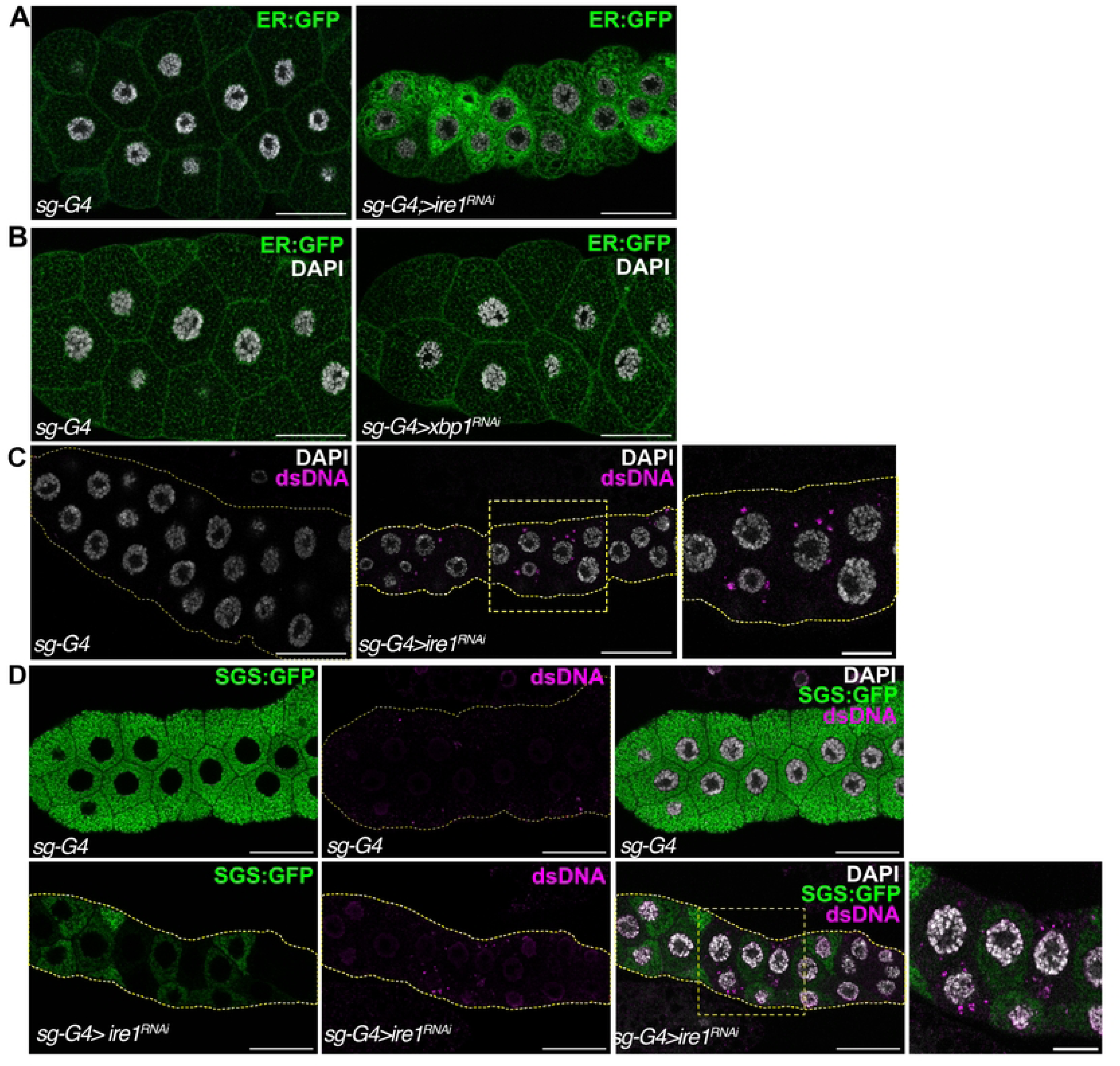
IRE-1 is required for proper ER development and preventing cytoDNA accumulation. (A) *Ire1* was depleted in the SG and their effect on the ER was visualized by ER-localized GFP (ER:GFP). A control SG (*sg-G4*) and a representative image of *ire1* depleted SGs are shown (*sg-G4>ire1^RNAi^*). (B) *Xbp1* was depleted in the SG (*sg-G4>xbp1^RNAi^*) and their effect on the ER network was visualized. (C) *Ire1* was depleted in the SG (*sg-G4>ire1^RNAi^*) and cytoDNA was visualized by anti-dsDNA (magenta). (D) The SGS:GFP genomic construct was used to monitor the expression of glue proteins (green) in control and *ire-1* depleted SGs at 110-120 hrs AEL. cytoDNA was also visualized by anti-dsDNA (magenta). (Scale bars for all the images: 50 μm and 20 μm for magnified images).

To further investigate the role of the IRE1 pathway, XBP1 or IRE1 was overexpressed in *de2f1b* SGs and determined if they can suppress the *de2f1b* SGs phenotypes. Interestingly, we discovered that overexpression of either factor had a dominant effect in wild-type SGs. Instead of having a mesh-like structure, overexpressing XBP1 or IRE1 resulted in SG cells in which the cytoplasmic space is filled with ER:GFP (Figure 6A, upper panel). Since many targets of XBP1 are important for ER biogenesis and function (11), XBP1 or IRE1 overexpression likely expands the ER compartment in the SG. However, overexpressing IRE1 had different effects in *de2f1b* SGs, while XBP1 overexpression produced a similar effect observed in the wild-type SG. Specifically, IRE1 overexpression in *de2f1b* SGs produced regions with intense ER:GFP signals that are scattered in the cytoplasm of *de2f1b* SG cells (Figure 6A, lower right panel). However, these cells maintained a mesh-like ER network and failed to expand ER compartment, as shown with XBP1 overexpression (Figure 6A magnified images). This result indicates that *de2f1b* mutations affect IRE1-dependent functions but not XBP1-dependent functions. Interestingly, in addition to the ER phenotype, we observed that the overexpression of either XBP1 or IRE1 in wild-type SGs was sufficient to produce weak yet discernible levels of cytoDNA (Figure 6B). Given that ER structures are abnormal in these cells (Figure 6A), this result, together with *ire1* depletion data (Figure 5), indicates that tight regulation of ER homeostasis is necessary for preventing cytoDNA accumulation during SG development.

**Figure 6.**
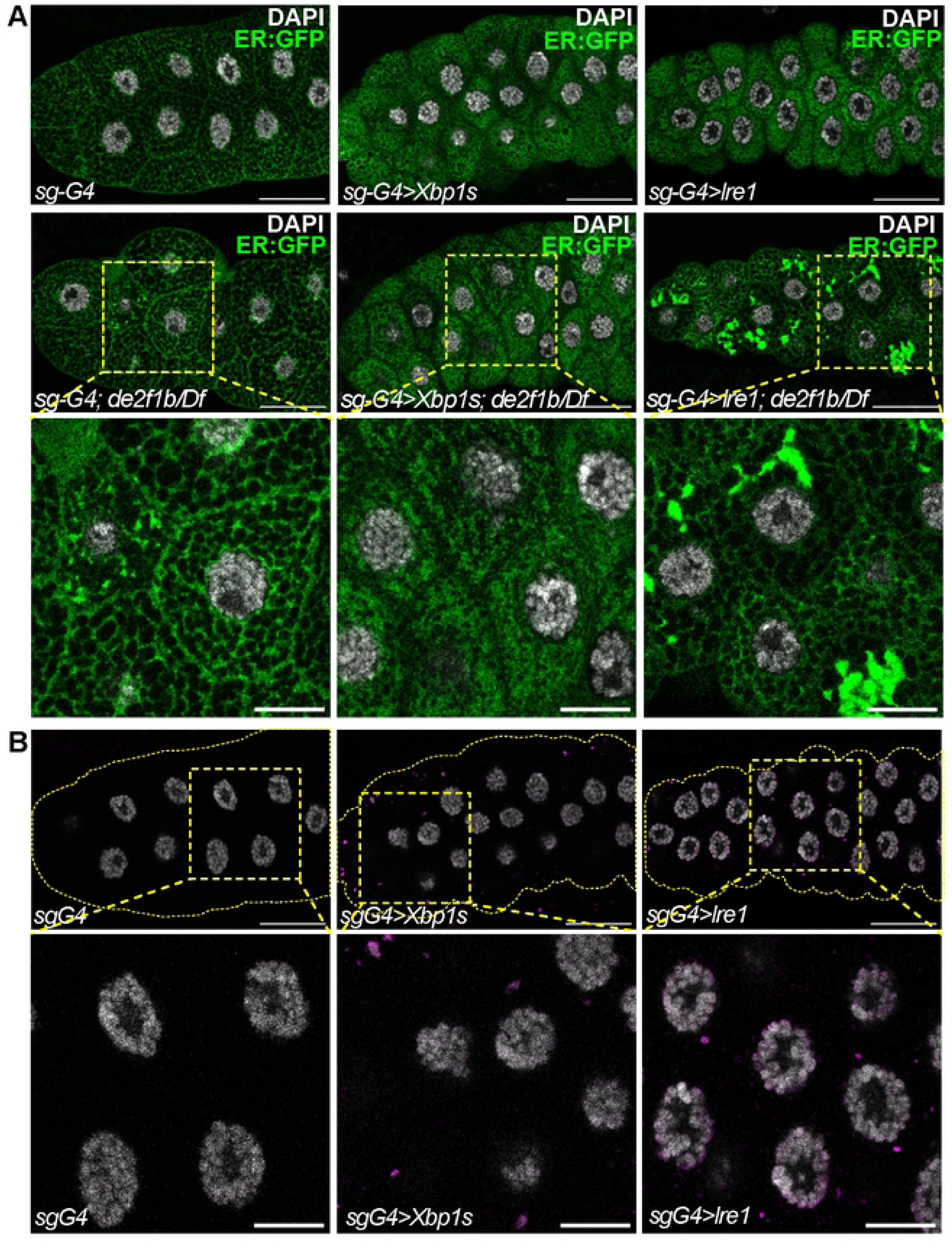
IRE1 overexpression differently affects the ER in *de2f1b* SGs. (A) XBP1 or IRE1 was overexpressed in control (*sg-G4*) and *de2f1b* SGs (*sg-G4; de2f1b/Df*), and the ER was visualized by ER:GFP. Magnified images of the boxed area are shown in the lower panels. (B) The effect of overexpressing XBP1 or IRE1 on cytoDNA accumulation in wild-type SGs is determined. The presence of cytoDNA was visualized by anti-dsDNA. Magnified images of the boxed area are shown in the lower panels. (Scale bars for all the images: 50 μm and 20 μm for magnified images).

## Discussion

The functional importance of E2F during endoreplication has been extensively studied in multiple model organisms (36–39). Due to its unique developmental characteristics, the fruit fly serves as an excellent model to investigate the role of E2F in endoreplication. In particular, the Drosophila larval salivary SG, which undergoes approximately 10 cycles of endoreplication, has provided significant insights into the regulatory function of E2F family proteins during endoreplication (40). Building on our previous findings that an alternatively spliced form of dE2F1, dE2F1b, is specifically required for endoreplicating tissues (16, 17), we now demonstrate that dE2F1b deficiency leads to heightened DNA damage, accumulation of cytoDNA, and disruption of ER homeostasis in the SG. Unexpectedly, we also discovered that the physiological activation of the IRE pathway is attenuated in *de2f1b* SGs, a process which is necessary not only for their fated role as a secretory tissue but also for preventing cytoDNA accumulation.

Mutations in *de2f1b* disrupt the negative feedback network that normally limits CycE/CDK2 activity, leading to deregulated S-phase entry during endoreplication (17). As a result, SG cells may initiate S phase without sufficient levels of proteins required for efficient DNA synthesis. This aberrant entry into S phase likely underlies the elevated DNA damage observed in *de2f1b* SGs (Figure 1A). The TUNEL assays (Figure 1B) also indicate that the DNA damage incurred in the absence of dE2F1b is not efficiently repaired. Given that numerous DNA repair genes are known to be regulated by E2F (3), the free DNA ends observed in *de2f1b* SG nuclei reflect impaired repair capacity. Importantly, the TUNEL-positive cells in *de2f1b* SGs are unlikely to be apoptotic, since transgene expression, such as ER:GFP, is maintained across all SG cells (Figure 3C), indicating that these cells remain biologically active. This is consistent with the known resistance of polyploid cells to apoptosis (41), which may allow SG cells to tolerate the accumulation of unrepaired DNA. Notably, previous studies have shown that excessive DNA damage and impaired DNA repair can lead to cytoDNA production (42). Therefore, cytoDNA observed in *de2f1b* SGs is probably of nuclear origin, accumulating over repeated cycles of endoreplication. This idea is further supported by the occasional presence of cytoDNA in the *de2f1b* larval fat body (Figure 3D), another endoreplicating polyploid tissue. However, we cannot rule out the contribution of mitochondrial DNA to cytoDNA accumulation, as E2F deregulation has also been linked to altered mitochondrial function (43). Isolation and sequencing the cytoDNA from *de2f1b* SGs will precisely determine the origin of cytoDNA.

CytoDNA is a well-established trigger of the IIR in mammals (44). Over the past decades, proteins that directly bind cytoDNA to elicit inflammatory responses, such as cGAS, have been identified and extensively studied (25). Upon binding cytoDNA, cGAS produces cyclic GMP-AMP (cGAMP), which is subsequently detected by STING at the ER, leading to activation of the NF-κB pathway (25, 45). While Drosophila possesses orthologs of both cGAS and STING, the functional counterpart to the mammalian cGAS that directly binds cytoDNA is yet to be identified (28). However, our findings show that the expressions of AMPs as well as Drosophila STING targets are upregulated in *de2f1b* SGs (Figure 2C and 2D). Interestingly, a previous study demonstrated that mutations in the lysosomal DNase, *DNase II*, can also trigger AMP expression (46). Although the precise role of the cGAS-STING pathway remains to be determined, it is plausible that cytoDNA is also a potent inducer of the inflammatory response in Drosophila.

It is interesting to note that the ER serves as a site for cytoDNA signaling and processing in mammals. The DNA exonuclease TREX1, which degrades cytoDNA, is localized to the ER, and STING is also an ER-resident protein, which translocates to the Golgi apparatus upon activation (47, 48). Importantly, while a Drosophila 3’-5’ DNA exonuclease, CG3165, is annotated as the ortholog of human TREX1, its role in cytoDNA processing remains uncharacterized. Nevertheless, any genetic manipulations that affected ER function in our study, such as *ire1* depletion and the overexpression of XBP1 or IRE1, resulted in cytoDNA accumulation (Figure 5C and Figure 6B). These results highlight the critical role of the ER in limiting cytoDNA during SG development. This may also explain the low but detectable levels of cytoDNA observed in SGs expressing SGS:GFP (Figure 3D). The replacement of the C-terminal sequence of the *SGS3* coding region with the GFP sequence may impair proper folding of the protein in the ER, potentially triggering ER stress and compromising the ER’s ability to efficiently eliminate cytoDNA. Indeed, expression of misfolding proteins has been used as a tool to induce the UPR in Drosophila (33).

Although the mechanism underlying ER’s role in cytoDNA elimination remains unclear, our data indicate that an XBP1-independent function of IRE1 is crucial for this process. Despite efficient depletion (S1 Fig), *xbp1* knockdown did not lead to ER defects or cytoDNA accumulation, while *ire1* depletion did (Figure 5). In addition to atypical splicing of *xbp1* mRNA, IRE1 degrades other targets through a process known as regulated IRE1-dependent decay (RIDD) (12, 35). Perhaps failure to degrade RIDD targets has a more substantial impact on maintaining ER homeostasis and limiting cytoDNA in the SGs. Supporting this notion, IRE1-dependent function seems to be primarily affected in *de2f1b* SGs while XBP1-dependent functions are not (Figure 6A). It will be important to investigate genes whose expressions are upregulated in *de2f1b* SGs, as they may represent the targets of RIDD that contribute to the *de2f1b* SG phenotype.

One of the most intriguing findings from our study is that the physiological activation of IRE1 is attenuated by E2F deregulation (Figures 4 and 6). This may explain why exocrine tissues are particularly vulnerable in *E2f1* and *E2f1/2* knockout mice (6–8). One plausible mechanism by which E2F affects the IRE1 pathway is that E2F directly or indirectly regulates the expression of genes crucial for IRE1 activity. Supporting this idea, RNA-seq analysis revealed that the expression of translocon subunits *sec61α* and *sec61β* are downregulated in *de2f1b* SGs (Figure 3B and Supplemental Table 1). The translocon complex facilitates the movement of newly synthesized proteins across the ER membrane and has been shown to play a critical role in IRE1 target specificity (49). By physically interacting with IRE1, the translocon complex positions IRE1 near its target RNA, including *xbp1,* at the ER membrane. In secretory tissues such as the SG, reduced levels of translocon components may impair specific molecular processes such as IRE1’s ability to recognize its targets. However, we cannot exclude the possibility that the decreased expression of the translocon subunits is the consequence and not the cause of IRE1 inhibition. Indeed, *ire1* or *xbp1* depletion resulted in a decrease in *sec61α* expression level (Figure 5C). An alternate mechanism by which dE2F1b deficiency attenuates IRE1 is that cytoDNA itself is the factor affecting IRE1 activity. Since cytoDNA elimination is an IRE1-dependent ER function, excessive production of cytoDNA during endoreplication of *de2f1b* SGs could lead to sustained ER stress. Previous studies have shown that chronic ER stress can attenuate the IRE1 pathway, mediated by other branches of the UPR (50, 51). Thus, it is conceivable that the prolonged ER stress in *de2f1b* SGs reduces IRE1 activity below its physiological level, disrupting ER network formation and impairing the ability to further process cytoDNA. We are currently investigating if other branches of the UPR, PERK and ATF6 pathways, are involved in IRE1 attenuation in *de2f1b* SGs.

If cytoDNA is the factor inhibiting the IRE1 pathway, perhaps cytoDNA serves as a signal for detecting abnormal SG cells during development. SG cells acquiring unrepairable DNA damage may accumulate excessive levels of cytoDNA during multiple rounds of endoreplication. This could attenuate IRE1 activity below its physiological level via chronic activation of the UPR, preventing the formation of the ER network necessary for glue protein production. This regulatory network would function as a safeguard mechanism to prevent SGs with unrepairable DNA damage from expressing defective glue proteins. Such a mechanism would be particularly beneficial for polyploid cells, as they cannot be easily eliminated by apoptosis (41). Interestingly, pancreatic cells in *E2f1/2* knockout mice become polyploid with age, correlating with the tissue atrophy (7). It will be interesting to investigate if those cells also accumulate cytoDNA and attenuate IRE1 activity. Overall, our findings provide crucial insights into why exocrine tissues are particularly susceptible to E2F deregulation and reveal a complex relationship between genome stability, cytoDNA accumulation, and ER homeostasis.

## Materials and methods

### Fly strains and culture

All Drosophila strains and genetic crosses were maintained at 25 °C on standard cornmeal medium. Complete genotypes of the flies used in each figure are provided in the S1 Table. The following flies were obtained from Bloomington Drosophila Stock Center (Bloomington, IN, USA): *Df(3R) Exel6186* (BDSC 7665), *Sgs3-GFP* (BDSC 5884), *UAS-mito-HA-GFP* (BDSC 8442), *UAS-Xbp1-EGFP. HG* (BDSC 60730), *UAS-Ire1^RNAi^* (BDSC 62156), *UAS-Xbp1^RNAi^* (BDSC 36755) and *Sgs3-GAL4* (BDSC 6870). The *de2f1b* mutant allele was previously described (16) and *UAS-XBP1s* and *UAS-IRE1* flies were generous gift from Dr. Hui-ying Lim at University of Alabama at Birmingham (52). Complete genotypes of the flies used in each figure are listed in S3 Table.

### Immunostaining

For immunostaining, wandering third instar larval (unless indicated otherwise) salivary glands were dissected in PBS and immediately fixed in 4% formaldehyde in PBS for 30 minutes at room temperature. Fixed tissues were then washed twice 10 minutes with 0.3% PBST (0.3% TritonX- 100 in 1XPBS) and once with 0.1% PBST (0.1% TritonX-100 in 1XPBS). Samples were incubated with primary antibodies in 0.1% PBST containing 5%NGS (normal goat serum) overnight at 4°C. Samples were then washed with 0.1% PBST, incubated with appropriate secondary antibodies in 0.1% PBST and 5% NGS for 2 hours at room temperature, followed by washes in 0.1% PBST prior to mounting. DNA was visualized with 0.1 μg/mL DAPI. Representative images were selected from a minimum of 10 independent tissues. The anti-𝛾H2Av (UNC93-5.2.1) and anti-double stranded dsDNA (autoanti-dsDNA) antibodies were obtained from the Developmental Studies Hybridoma Bank (https://dshb.biology.uiowa.edu). For anti-dsDNA immunostaining, confocal microscopy parameters were optimized to minimize mitochondrial signal detection while preserving visualization of cytoDNA signals.

### Microscopy and image processing

All fluorescently labeled tissues were mounted using a glycerol-based anti-fade mounting medium containing 5% N-propyl gallate. Images were acquired using a laser-scanning Leica SP8 confocal microscope using either 40x/1.3 oil immersion objective or 63x/1.4 oil immersion objectives at the Advanced BioImaging Facility, McGill University. Representative images are individual slices from z-stacks. All images were processed using Fiji (http://fiji.sc/Fiji).

### Gene expression Analysis

For RT-qPCR, RNA was extracted from salivary glands from L3 larvae of the appropriate genotype using the RNeasy Mini Kit (Qiagen #74104). To eliminate genomic contamination, the RNA was treated with RNase-free DNase I. 300 ng of RNA was used to synthesize cDNA using the iScript™ cDNA Synthesis Kit (Bio-Rad #1708891). To analyze the gene expression qPCR was performed using DyNamo Flash SYBR Green qPCR kit (Thermo Scientific™ # F415XL) according to the manufacturer’s specification. The primers used in this study are mentioned in S2 Table.

### Quantification of the 𝛾-H2Av foci

To distinguish the 𝛾-H2AV foci from the random background noise, the Difference of Gaussian (DoG) algorithm was applied to the confocal images using the Gaussian blur filter in Fiji. By subtracting duplicate images at different blur strengths (image1 (σ=1) - image2 (σ=2)), the high-frequency spatial information, including the background noise, was removed. Thresholding was then used to preserve only the high-intensity signals that are the 𝛾-H2AV foci. The number of pH2AV foci was counted for every nucleus in a focal plane using the Analyze Particles function.

### TUNEL assay

For the TUNEL assay, tissues were dissected in 1X PBS and fixed in 4% formaldehyde in PBS for 20 minutes at room temperature. Fixed samples were washed with 2% Triton for 1 hour at RT and TUNEL assays were performed using In Situ Cell Death Detection Kit TMR red (Roche #12156792910) according to the manufacturer’s specification.

### RNA sequencing and bioinformatics

RNA sequencing was conducted on an Illumina NovaSeq 6000 platform to generate paired-end reads of 100 bp with 25M reads per sample. The quality of the raw reads was assessed with FASTQC v0.11.8. After examining the quality of the raw reads, no trimming was deemed necessary. The reads were aligned to the fly reference genome with STAR v2.7.6a, with a mean of 87 % of reads uniquely mapped. The raw counts were calculated with FeatureCounts v1.6.0 based on the fly reference genome (release 102). Differential expression was performed using the DESeq2 R package and differentially expressed genes were defined as those with an adjusted p-value < 0.05 (Benjamini–Hochberg correction) and a log2 fold change threshold of ±1.

All raw and processed RNA-seq data have been deposited in GEO under accession number GSE294319. Gene ontology (GO) and pathway enrichment analyses were performed using ShinyGO V0.81using the KEGG pathway database (http://bioinformatics.sdstate.edu/go/) Ge SX, Jung D & Yao R, Bioinformatics 36:2628–2629, 2020. In this result, the most significant (lowest FDR) downregulated genes were categorized as "Protein processing in the endoplasmic reticulum". The gene ontology figure showing the top 5 most significant downregulated KEGG pathways was generated in R 4.3.3 using ggplot2 (v3.5.; Wickham H, 2016). The complete list of downregulated genes in *de2f1b* mutant salivary glands and gene ontology analysis result are provided in the S1 Table.

### Quantification of XBP1 splicing

Mean intensity for DAPI and XBP1-GFP was measured for each nucleus in focus by manually selecting each nucleus in Fiji. The mean intensity of the XBP1-GFP signal was divided by the mean intensity of DAPI signal for each nucleus.

### Statistical analysis

Two-tailed unpaired t-test was used for all the RT-qPCR results. For nuclear intensity ratio of DAPI and XBP1-GFP intensity and 𝛾-H2Av quantification two-tailed Mann-Whitney test was used. For all statistical analyses, p<0.05 was considered statistically significant. p-values represent ns = *p* > 0.05; * = *p* < 0.05; ** = *p* < 0.01; *** = *p* < 0.001, **** = *p* < 0.0001.

## Acknowledgements

The authors are grateful to Dr. Hui-Ying Li (University of Alabama at Birmingham) for sharing *UAS-Ire1* and *UAS-*Xbp1 fly stocks. The authors also thank Bloomington Drosophila Stock Center (NIH P40OD018537) for fly stocks used in this study. We thank the IRCM Bioinformatics core facility for their support on bioinformatics analysis.

## Supplemental information captions

**S1 Fig.** Knockdown efficiency of *ire1^RNAi^* and *xbp1^RNAi^* constructs in the salivary glands using *Sgs3-GAL4*.

**S1 Table:** List of downregulated genes in *de2f1b* mutant salivary gland from RNA-seq data and results of gene ontology analysis on downregulated genes.

**S2 Table:** Complete genotypes of the Drosophila stocks used in each figure.

**S3 Table:** Primer sequences used in this article for qPCR.

**S1 Data:** Summary statistics and statistical tests used for all figure panels.

